# BioLLMNet: Enhancing RNA-Interaction Prediction with a Specialized Cross-LLM Transformation Network

**DOI:** 10.1101/2024.10.02.616044

**Authors:** Md Toki Tahmid, Abrar Rahman Abir, Md. Shamsuzzoha Bayzid

**Affiliations:** Bangladesh University of Engineering and Technology

## Abstract

Existing computational methods for the prediction of RNA related interactions often rely heavily on manually crafted features. Language model features for bio-sequences has gain significant popularity in proteomics and genomics. However, during interaction prediction, how language model features from different modalities should be combined to extract the most representative features is yet to be explored. We introduce BioLLMNet, a novel framework that introduces an effective combination approach for multi-modal bio-sequences. BioLLMNet provides a way to transform feature space of different molecule’s language model features and uses learnable gating mechanism to effectively fuse features. Rigorous evaluations show that BioLLMNet achieves state-of-the-art performance in RNA-protein, RNA-small molecule, and RNA-RNA interactions, outperforming existing methods in RNA-associated interaction prediction.

## 1 Introduction

Ribonucleic acids (RNAs) are essential biomolecules that play diverse roles in cellular processes through interactions with other RNAs, proteins, and small molecules [20, 18, 3, 9, 24, 33, 8, 31]. Understanding these RNA-associated interactions is crucial, as they regulate various physiological and pathological processes. Specifically, RNA-RNA interactions are involved in post-transcriptional processes, contributing to gene expression regulation [26, 12]. RNA-protein interactions are vital for maintaining cellular homeostasis, and disruptions in these interactions can lead to cellular dysfunctions or diseases such as cancer [29, 1, 15, 6]. Furthermore, RNA-small molecule interactions have significant implications in therapeutic development, as RNAs can serve as potential drug targets, especially when conventional protein targets are less accessible [5, 16].

Despite the importance of these interactions, the discovery of RNA-associated interactions is often challenging due to the complexity of RNA structures and the limited availability of experimentally resolved RNA-containing structures [17, 11]. Existing computational methods have therefore been developed to predict such interactions and facilitate research in this area. These approaches typically involve encoding interacting molecules into computer-recognizable features such as sequences, structures, and physicochemical properties, followed by integrating these features to predict interactions. For RNA-RNA interactions, tools like MD-MLI and lncIBTP convert RNA sequences and secondary structures into numerical vectors, then use machine learning algorithms to identify potential interaction partners [25, 32]. In the case of RNA-protein interactions, tools such as CatRAPID and PRPI–SC use both sequence and structural information to calculate interaction scores, combining features like hydrogen bonding potential, charge, and hydrophobicity [34, 2]. For RNA-small molecule interactions, LigandRNA and dSPRINT extract molecular fingerprints and physicochemical properties of small molecules, then predict binding affinities with RNA using statistical models and similarity-based ranking [23, 7].

While existing methods have made significant progress, they face several limitations. Most approaches rely on manually crafted features based on sequence motifs, structural properties, or physicochemical characteristics, which may not fully capture the complexity and dynamic nature of RNA interactions. Additionally, these methods often lack flexibility in adapting to diverse RNA sequences and their interaction partners. Notably, none of the current models utilizes RNA language model features, which have shown great potential for capturing intricate sequence patterns and contextual information in a data-driven manner. Integrating RNA language model features could provide a more comprehensive understanding of RNA interactions, enhancing the predictive power and generalizability of these methods. Therefore, there is a need for methods that combine language model representations to more accurately predict RNA-RNA, RNA-protein, and RNA-small molecule interactions.

To address these issues, we propose a novel framework **BioLLMNet** for predicting RNA interactions with protein, RNA and small molecules. To the best of our knowledge, we are the first to incorporate RNA language model embeddings for predicting RNA-associated interactions, incorporating advanced sequence representations that capture complex RNA patterns. Moreover, We employ language model embeddings for both modalities in interaction prediction, meaning that for RNA and its interacting partners (whether proteins, small molecules, or other RNAs), we utilize language model-based representations. This approach allows for a more comprehensive and context-aware understanding of interaction patterns. Additionally, for biological language model feature fusion for different modalities, we propose a learnable gating mechanism that adaptively combines features from both RNA and the other modality. Finally, through rigorous evaluation we validate that BioLLMNet achieves state-of-the-art performance for predicting RNA-RNA, RNA-protein, and RNA-small molecule interactions, outperforming existing approaches in all categories.

## 2 Methods

In this section, we present the architecture of BioLLMNet, focusing on embedding processes for RNA, protein, and small molecules, as well as the transformation and gated combination of multimodal feature spaces. The overall architecture of BioLLMNet is shown in Figure 1.

**Figure 1.**
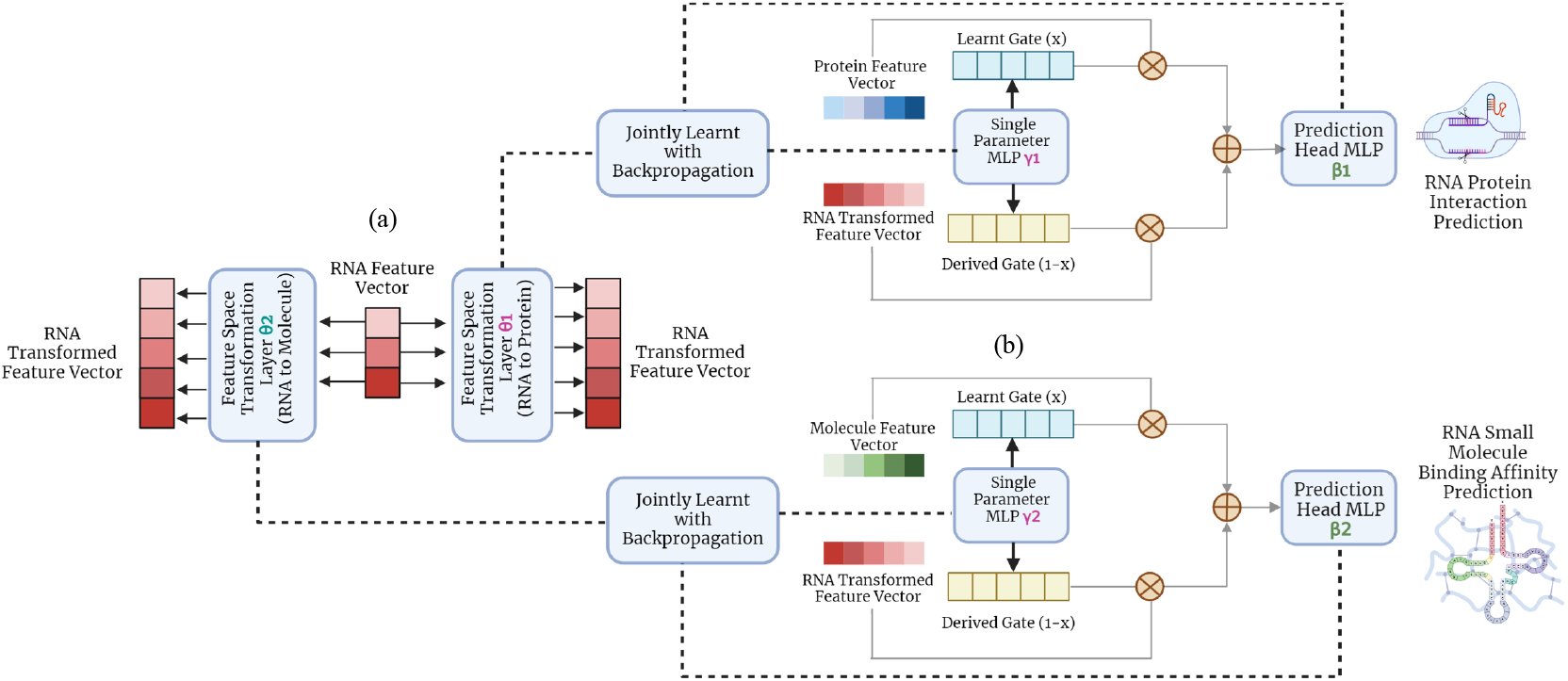
Overall architecture of BioLLMNet. (a) We transform the feature space size of RNA to match the feature space size of the protein or molecules. For that, we employ a single layer MLP which performs this transformation. This MLP is learned with the whole network. (b) Finally, we use a gated weight to first perform a weighted average of the two sequence embeddings (which are now in the same dimension after step 2), and then pass the joint embedding through a multi-layer MLP network which is essentially the prediction head. These three MLPs (transformation, gated weight, and prediction head) are learned jointly with backpropagation.

### RNA Language Model Representation

RNA sequences composed of adenine (A), uracil (U), cytosine (C), and guanine (G) are embedded using RNA-FM, which converts sequences 𝒮 = {*s*_1_, *s*_2_, …, *s*_*n*_} into high-dimensional feature vectors for downstream tasks. We use BiRNA-BERT[**?**] to generate language model features for RNA sequences.

### Protein Language Model Representation

Protein sequences 𝒫 = *p*_1_, *p*_2_, …, *p*_*m*_ are embedded using esm2 [], which tokenizes the sequence and processes it into a 1024-dimensional space using transformer layers, with the CLS token capturing a global representation.

### Molecular Language Model Representation

MoleBERT [30] tokenizes SMILES strings *S* by converting them into molecular graphs *G* = (*V, E*). Each atom *v*_*i*_ is encoded into a latent vector *z*_*i*_, quantized into a token *t*_*i*_, and embedded into a feature vector *e*_*i*_. The final set of embeddings *E* = {*e*_1_, *e*_2_, …, *e*_*n*_} captures the molecule’s structural context.

### 2.1 Transformation of Cross-Lingual Feature Space

To unify the multi-lingual biological feature space, we transform lower-dimensional modality vectors into higher-dimensional ones using Algorithm 1. For RNA-protein and RNA-small molecule interactions, we use a single-layer MLP to map the RNA features to the dimensionality of the other modality, followed by a ReLU activation. This transformation ensures consistent dimensionality for downstream tasks and is learned during training.

#### Algorithm 1

Cross-Lingual Feature Transformation

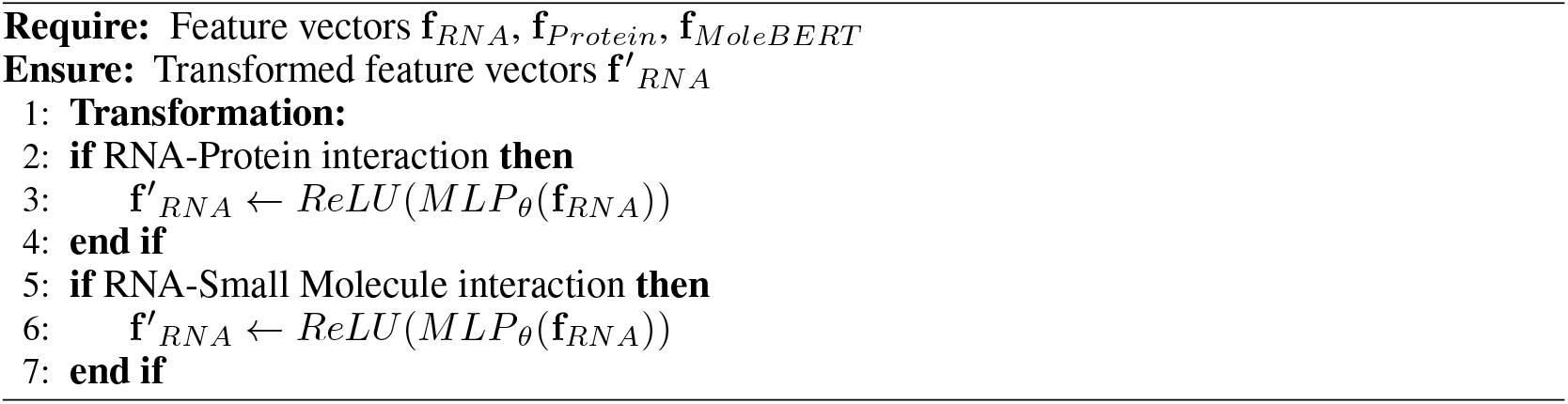

### 2.2 Gated Combination of Transformed Spaces

After transforming the feature spaces to the same dimensionality, we combine them using a gated mechanism with Algorithm 2. This approach dynamically balances the contribution of each modality by learning a gate for each feature dimension. The gate allows the model to emphasize one modality over another based on input data, improving feature relevance for downstream tasks.

The gate parameters are learned via backpropagation, and the final prediction is made through a three-layer deep neural network, optimized with a combined loss function.

## 3 Results

### RNA Protein Interaction Prediction

For the RNA-protein interaction prediction, we evaluate BioLLMNet on the RPI-1460 dataset, comparing it with six models: RPISeq-RF, IMPIner, CFRP, RPITER, LPI-CSFFR, and RNAincoder. From Table 1 we see that, BioLLMNet achieves superior performance across all metrics, showing an 11.6% improvement in MCC, 4.8% in accuracy, 9.0% in F1, and 9.6% in precision on RPI-1460.

**Table 1:** Performance comparison of different methods on RPI1460 dataset.

| Method | MCC | ACC | F1 | Precision | Recall |
| --- | --- | --- | --- | --- | --- |
| RPISeq-RF[19] | 0.570 | 0.780 | 0.780 | 0.790 | 0.780 |
| IMPIner[21] | 0.520 | 0.760 | 0.770 | 0.720 | 0.830 |
| CFRP [4] | 0.630 | 0.810 | 0.820 | 0.830 | 0.780 |
| RPITER [22] | 0.412 | 0.690 | 0.510 | 0.610 | 0.480 |
| LPI-CSFFR [10] | 0.600 | 0.830 | 0.840 | 0.780 | 0.910 |
| RNAincoder [27] | 0.760 | 0.880 | 0.840 | 0.810 | 0.940 |
| BioLLMNet | **0.848** {↑11.6%} | **0.923** {↑4.8%} | **0.925** {↑9.0%} | **0.888** {↑9.6%} | **0.966** {↑2.8%} |

#### Algorithm 2

Gated Combination of Transformed Spaces

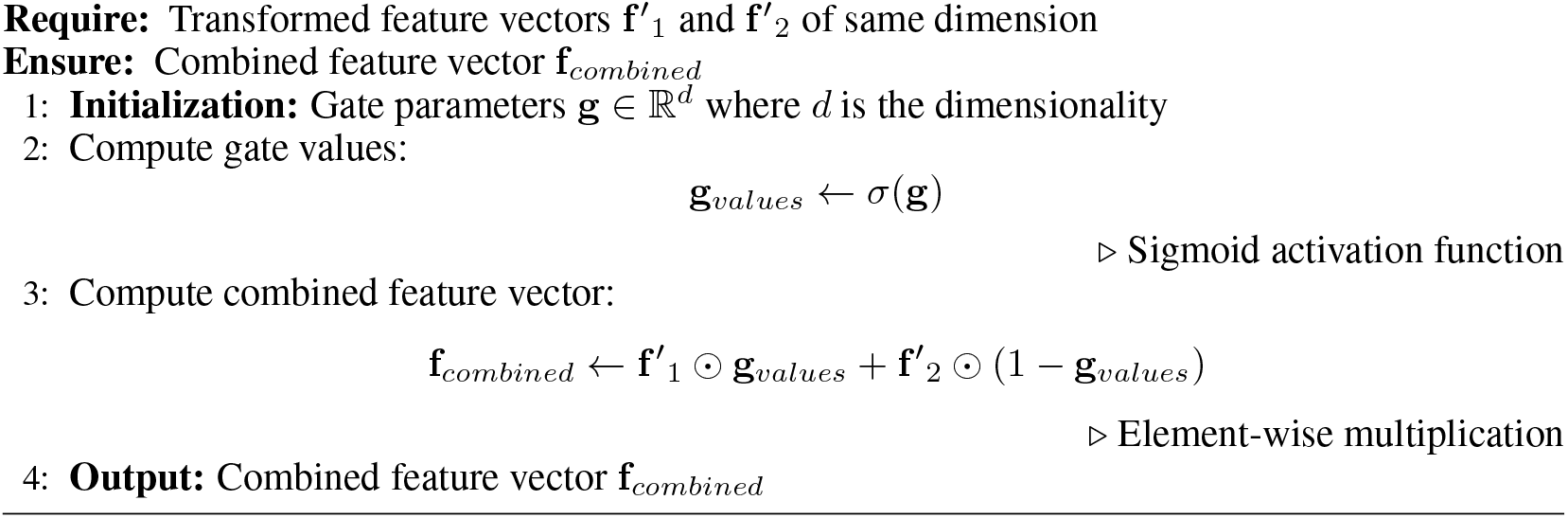

### Case Study

We conducted a case study to evaluate BioLLMNet’s ability to identify RNA-protein interactions, focusing on the RNA sequence ‘2QEX-0’, which interacts with 37 different proteins. As shown in Figure 2, BioLLMNet correctly predicts all interactions with proteins from the 2QEX family (29 proteins) but makes two incorrect predictions with 4GD1-G and 2WH1-Y. Upon examining the confidence scores, the incorrect predictions show significantly lower scores, demonstrating BioLLMNet’s robustness with 100

### RNA-Small Molecule Interaction Prediction

RNA-small molecule interaction prediction is a challenging regression task where given a RNA sequence and molecular representation of a small moleule we need to determine their binding affinity. For that, we use the benchmarking datasets provided in https://web.iitm.ac.in/bioinfo2/RSAPred/Predict.html where RNA sequences of six RNA families : miRNA, Viral RNA, Riboswitch RNA, Ribosomal RNA, Repeats RNA and Aptamers are provided along with the proteins they interact with. Following RSAPred [], we divide the dataset into training and validation set with 20% validation split and report the pearson correlation and the mae scores in Table 2. We see that, BioLLMNet achieves superior performance in five out of the six datasets in pearson’s correlation and achieves best mae score in all datasets.

**Table 2:**
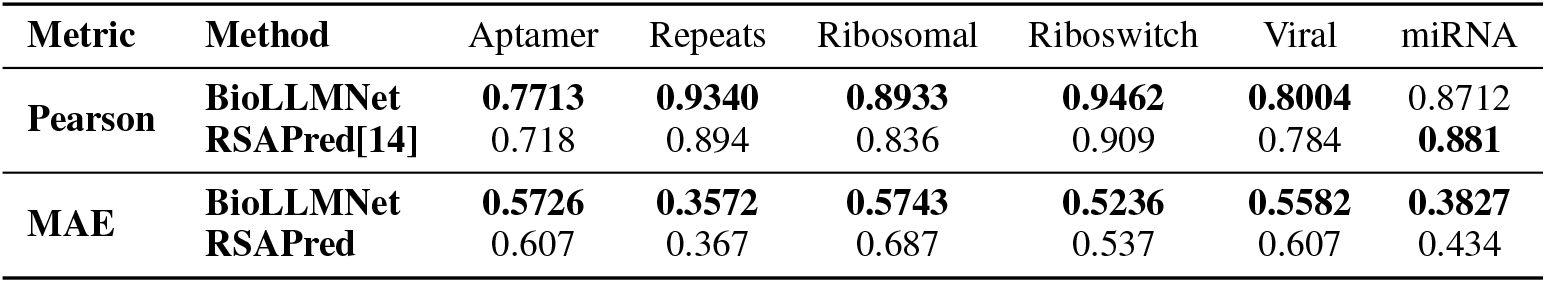
Comparison of BioLLMNet and RSAPred on various datasets (Pearson and MAE).

| Metric | Method | Aptamer | Repeats | Ribosomal | Riboswitch | Viral | miRNA |
| --- | --- | --- | --- | --- | --- | --- | --- |
| Pearson | BioLLMNet | **0.7713** | **0.9340** | **0.8933** | **0.9462** | **0.8004** | 0.8712 |
|  | RSAPred[14] | 0.718 | 0.894 | 0.836 | 0.909 | 0.784 | **0.881** |
| MAE | BioLLMNet | **0.5726** | **0.3572** | **0.5743** | **0.5236** | **0.5582** | **0.3827** |
|  | RSAPred | 0.607 | 0.367 | 0.687 | 0.537 | 0.607 | 0.434 |

**Figure 2.**
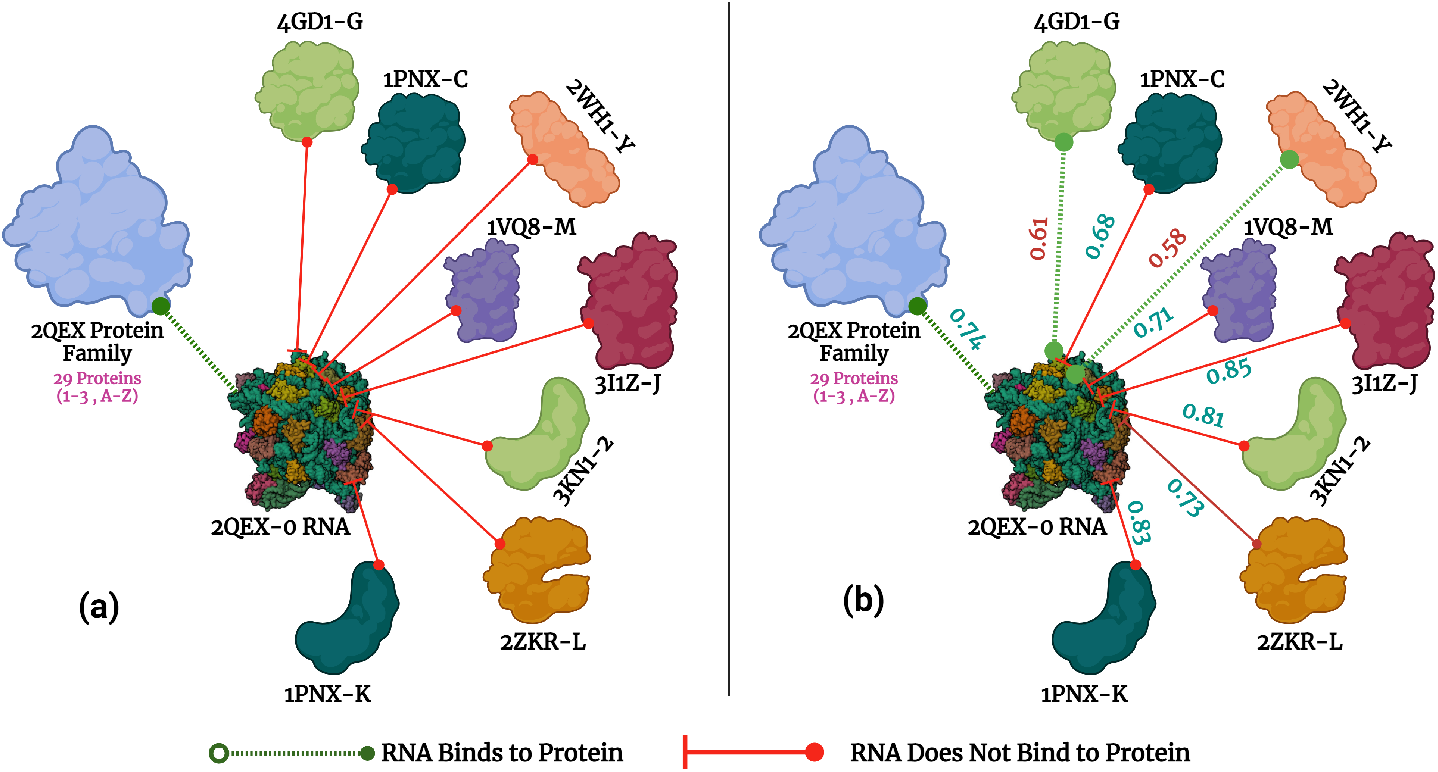
Case study of interactions of the 2QEX-0 RNA complex with different protein complexes. To the left (a), there is the actual interactions. 2QEX-0 interacts will all 29 proteins from the 2QEX protein family however, does not interact with the other eight proteins. (b) Predicted interaction result with BioLLMNet. BioLLMNet predicts the true interactions perfectly, however it predicts two non-interacting edges as interacting edges.

### Transformation Along Same Language– lncRNA-miRNA Interaction Prediction

The core idea of BioLLMNet is combining sequences from different modalities, requiring feature space transformation. We explore whether gated combinations work for the same modality by applying it to miRNA-lncRNA interaction prediction, where both sequences belong to the RNA domain but differ significantly in length. We use three RNA-RNA interaction datasets for evaluation with cross-validation across six train-test combinations. miRNA sequences range from 10-50 nucleotides, while lncRNA sequences vary from 200-4000 nucleotides. BioLLMNet outperforms CORAIN in four out of six dataset combinations (ATH-GMA, GMA-ATH, MTR-ATH, MTR-GMA), with up to 17.2% improvement as shown in Table 3. In ATH-MTR, BioLLMNet achieves 72% accuracy, and in GMA-MTR, it records 84% accuracy, with a 9.7% performance drop compared to the SOTA.

**Table 3:** Comparison of BioLLMNet, CORAIN, and PMLIPred on miRNA-lncRNA interaction datasets.

| Model | Train-Test Dataset |  |  |  |  |  |
| --- | --- | --- | --- | --- | --- | --- |
|  | ATH-GMA | ATH-MTR | GMA-ATH | GMA-MTR | MTR-ATH | MTR-GMA |
| PMLIPred[13] | 65 | 70 | 55 | 87 | 53 | 71 |
| CORAIN[28] | 69 | **74** | 67 | **93** | 58 | 84 |
| BioLLMNet | **69** {**↑0.0%**} | 72 {**↓2.7%**} | **69** {**↑3.0%**} | 84 {**↓9.7%**} | **68** {**↑17.2%**} | **85** {**↑1.2%**} |

## 4 Conclusion

In conclusion, BioLLMNet introduces an innovative approach to combining language model features from different bio-sequence modalities, providing a robust solution for RNA-associated interaction prediction. By transforming the feature space and employing a learnable gating mechanism, BioLLMNet effectively fuses multi-modal features, leading to state-of-the-art performance across various interaction types, including RNA-protein, RNA-small molecule, and RNA-RNA interactions.

Looking ahead, the framework’s potential can be further expanded by incorporating additional modalities such as structural or physiochemical features, which can be integrated in a similar manner to enhance prediction accuracy. Furthermore, future extensions could adopt strategies akin to LLAVA, where joint embeddings from natural language and visual modalities are learned, enabling the seamless incorporation of diverse feature types such as 3D molecular structures and other high-dimensional bio-sequence features. This will allow for the development of more sophisticated models capable of leveraging heterogeneous data sources to improve the accuracy and robustness of RNA-related interaction predictions.

